# Data-driven models of optimal chromatic coding in the outer retina

**DOI:** 10.1101/2022.02.07.479405

**Authors:** Luisa Ramirez, Ronald Dickman

**Affiliations:** Departamento de Física, Universidade Federal de Minas Gerais, CEP 31270–901 Belo Horizonte, Minas Gerais, Brasil

## Abstract

The functional properties of the outermost retinal circuits involved in color discrimination are not well understood. Recent experimental work on zebrafish has elucidated the in-vivo activity of photoreceptors and horizontal cells as a function of the stimulus spectrum, highlighting the appearance of chromatic-opponent signals at the first synaptic connection between cones and horizontal cells. These findings, together with the observed lack of gap junctions, suggest that the mechanism yielding early color-opponency in zebrafish is dominated by inhibitory feedback. We discuss the observed neuronal activity in the context of efficient codification of chromatic information, hypothesizing that opponent chromatic signals provide optimal codification, minimizing signal redundancy. We examine whether these functional properties are general across species by studying the dynamic properties of dichromatic and trichromatic outer retinal networks. Our findings show that dominant inhibitory feedback mechanisms provide an unambiguous codification of chromatic stimuli, whereas this property is not guaranteed in networks with strong excitatory inter-cone connections, for example via gap junctions. This provides a plausible explanation for the absence of gap junctions observed in the outermost zebrafish retinal layers. In addition, our study suggests that the simplest zebrafish-like network with dominant inhibitory feedback capable of optimally codifying chromatic information requires at least two successive inhibitory feedback layers. Finally, we contrast the chromatic codification performance of zebrafish-inspired retinal networks with networks having different opsin combinations. We find that optimal combinations lead to a chromatic codification improvement of only 13% compared with zebrafish opsins, suggesting that the zebrafish retina performs nearly optimal codification of chromatic information in its habitat.

**Author summary:** Recent experimental work has evidenced that outer neuronal circuits in the zebrafish retina use color-opponent mechanisms to codify and transmit chromatic information at the first synaptic contact between cones and horizontal cells. Inspired by these findings, we propose a data-driven model to study physiological and dynamical properties of outer retinal networks and their implications for color codification across vertebrate retinal circuits. We first study our model in a large parameter space, finding that the primary biological mechanism leading to color-opponent signals is mediated by dominant inhibitory feedback, e.g., via horizontal cell synaptic connections. In contrast, strong coupling among cones leads to ambiguous chromatic codification, undesirable in the outer retina. Then, we parameterize the model using zebrafish experimental data and quantify its chromatic codification performance. Our results suggest that trichromatic retinas with inhibitory feedback are highly efficient and capture most of the chromatic information variance typical from zebrafish environments. More specifically, a comparison among zebrafish-inspired retinal networks suggests that zebrafish retinal circuits are near-optimal chromatic codification of their natural chromatic information.

## 3 Introduction

The study of how chromatic information is processed in the retina has a long history [1–4]. Nevertheless, some questions on the specific neuronal circuits and their functionality remain unanswered[2, 5]. One approach that has provided insights into these questions is the study of visual systems from an information theory perspective, in which efficient information transfer plays a key role [6]. In retinal color processing, optimization is associated with reliable and efficient chromatic stimulus codification and transmission via neuronal responses and synaptic interactions [7–9]. Optimal codification, from this viewpoint, maximizes the information transfer rate in the limited retinal channel capacity [6, 10, 11].

Retinal circuits are organized in layers, such that the outermost neurons receive photonic stimuli, while the innermost neurons connect to downstream brain circuits. Photoreceptor neurons, specifically cones, are the basis of retinal color response in the outer retinal layer. They are categorized by their independent response to chromatic stimuli via the sensitivity function, which displays a vast diversity across species[3]. In retinas, photoreceptor types and their retinal connectivity determine the species-specific color space [3, 9]. For instance, human retinas have three types of photoreceptors, sensitive to long-, middle-, and short-wavelength stimuli, that shape a trichromatic visual system. Responses from this outermost layer activate horizontal cells (HCs), which are the next neurons in the retinal pathway. Similarly to photoreceptors, HC diversity across species is vast but their functionality and importance in chromatic retinal circuits is not fully understood [12]. HCs connect to several photoreceptors, providing lateral connectivity and consequently, integration of spatially localized responses. Signals from HCs are further processed by inner retinal layers and then transmitted to downstream neuronal circuits.

While the selective advantage conferred by color vision derives from object discrimination via spectral contrast (as opposed to mere brightness contrast), the response curves of L, M and S opsins have significant overlap, yielding highly redundant signals among the cone types. From the viewpoint of information theory, reducing such redundancy optimizes stimulus codification and the information transfer rate[13]. More specifically, theoretical work by Buchsbaum and Gottschalk [11] on trichromatic visual systems shows that given the covariance matrix of the photoreceptor responses, the eigenvalue or principal component transformation is optimal at decorrelating the chromatic information of these responses, resulting in efficient information transmission. Such a transformation leads to the appearance of chromatic opponent signals, also predicted by color-opponency theory [14]. This implies that photoreceptor responses turn inhibitory in certain ranges of the spectrum, and, are excitatory otherwise, guaranteeing their linear independence. Over the past decades, experimental work [15, 16] has provided evidence of such opponent responses to different spectral stimuli in retinal-ganglion layers and downstream neuronal circuits in vertebrates. Evidence of opponency in the outermost retinal circuits, however, is more recent [17, 18], and suggests that photoreceptor chromatic responses turn opponent at the first synapse with HCs.

The experimental works on the outermost retinal layers measure the in-vivo response of photoreceptors to different chromatic stimuli, including the response of horizontal cells. Specifically, Yoshimatsu et al. [17] found experimental evidence of chromatic-opponent responses in the outermost retinal circuits of zebrafish, yielding important insights into retinal color processing. Their work highlights the importance of horizontal cells in color codification and information transmission, and suggests that opponent cone responses depend on inhibitory feedback from HCs, whereas excitatory inter-cone connections via gap junctions are negligible. Thus, efficient coding of the chromatic information available in the typical zebrafish environment is attained at the earliest stage of visual processing. This raises two questions: First, whether these observations can be extrapolated to other visual systems, and second, whether optimizing codification and transference of chromatic information is the reason behind the observed neuronal behavior.

In this work, we study different outer retinal network architectures using a neuronal dynamic model based on observed zebrafish retinas [19]. We search for general properties that can be attributed to a broad set of species, such as the roles of excitatory and inhibitory synaptic connections. Although some works [20, 21] report evidence of gap junctions between cones in some vertebrates, their functional purpose in chromatic circuits remains unclear. We study whether the absence of gap junctions (as observed in zebrafish) facilitates coding of chromatic information, implying that such inter-cone connections, if present, are not crucial in color discrimination. Second, we investigate zebrafish-like network architectures, quantifying their performance in the context of optimal codification and information transfer, given the chromatic information typical of the zebrafish environment [22]. Specifically, we investigate the minimal architecture, including interneurons, optimizing codification of the available chromatic information. Some experiments have shown that some invertebrate species, such as butterflies, exhibit early chromatic opponency without interneurons [23]. Nevertheless, in this work, we investigate only retinal circuits that include interneuronal connections to yield opponent responses, as is the case in zebrafish.

## 4 Results

### 4.1 Dynamics of outer retinal networks

We begin by studying the dynamic properties of dichromatic and trichromatic networks with photoreceptors sensitive to short (B), middle (G) and long (R) spectral ranges. Figure 1a sketches a typical dichromatic network with inhibitory feedback, that is, with inhibitory HC-cone synaptic connections. More precisely, light stimuli induce linear excitatory responses, *I*_*i*_, of photoreceptor neurons (R and G), which in turn activate horizontal cells (H). The integrated response of horizontal cells, *I*_*H*_, provides an inhibitory feedback to photoreceptor neurons, inducing final responses, 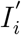, sent to downstream retinal circuits. Networks with excitatory intercone connections and inhibitory feedback have additional cone-cone connections (dashed arrow in Fig. 1a). We denote networks with dominant inhibitory feedback and weak inter-cone connections as Type I; networks with strong excitatory inter-cone connections are denoted Type II.

**Figure 1:**
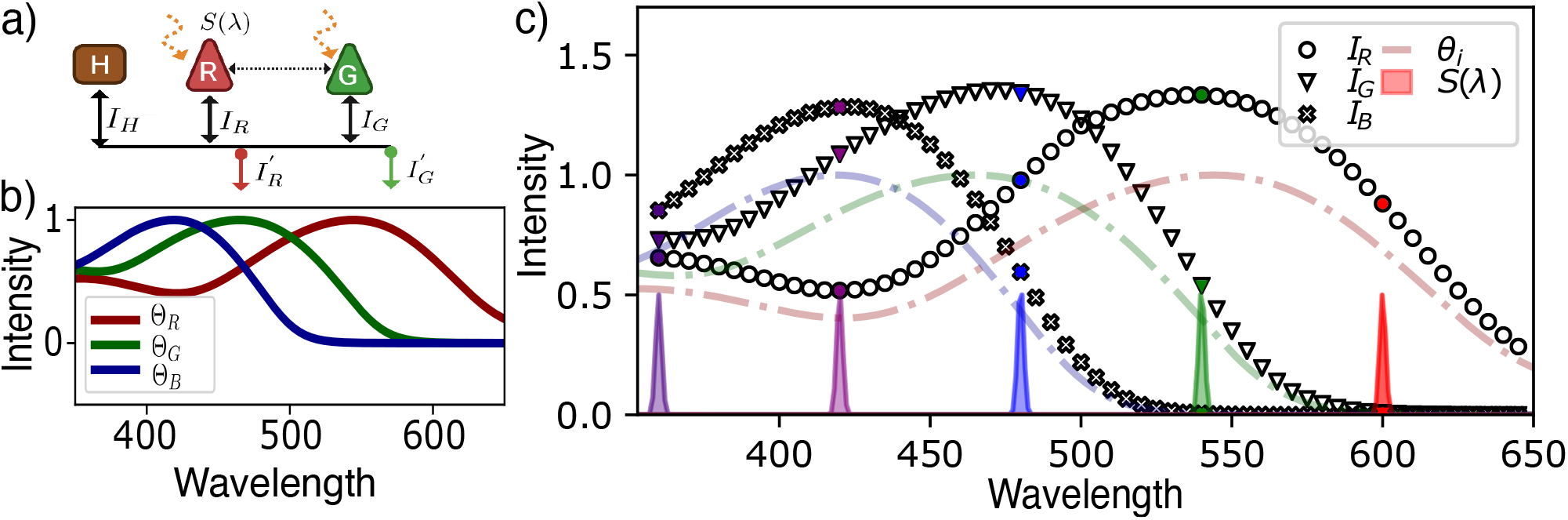
a) Dichromatic sketch of a fully connected network with an external stimulus, *S*(*λ*). R: red cones, G: green cones, B:blue cones, H: horizontal cells. Solid black arrows represent inhibitory synaptic conections on cones from horizontal cells. Dashed black arrows represent excitatory synaptic connections between cones. b) Sensitivity functions of independent red, green and blue zebrafish opsins [24]. c) Independent responses of photoreceptors to narrow gaussian stimuli. Dashed curves are red, green and blue zebrafish sensitivity curves and colored distributions correspond to five Gaussian stimuli normalized to exhibit a maximum intensity of 0.5, with variance *σ* = 1nm. Markers correspond to the independent responses, described by Eq. 1, of the three photoreceptors. Colored markers show the response to the five plotted stimuli.

One way of characterizing visual stimuli is via the spectral density, *S*(*λ*), which contains all relevant chromatic information. To calculate the isolated cone response, *I*_*i*_, to a stimulus with spectral density, *S*(*λ*), we integrate the product of Θ_*i*_(*λ*), the corresponding sensitivity function, and *S*(*λ*), over the spectrum:

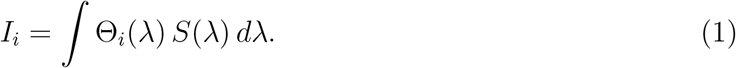

Determining responses to specific wavelength intervals requires stimuli with a narrow spectral distribution, conveniently represented by Gaussian distributions centered at a characteristic wavelength *λ*_0_, and having a small standard deviation; we use *σ*_0_ = 1nm in the present analysis. Photoreceptor sensitivity functions, on the other hand, can be broadly categorized into ultraviolet, blue, green and red according to the spectral range at which their response is maximum. Fig. 1b. shows the sensitivity functions of red, green and blue zebrafish cones[24], which we use as a template for our analysis. These curves were obtained experimentally using LEDs ranging from 350nm to 650nm, with the same luminance [17]. With these sensitivity functions and the Gaussian chromatic stimuli previously described, we use Eq. (1) to model photoreceptor responses. As shown in Fig. 1c, Gaussian chromatic stimuli with the same luminance intensity yield responses similar to the sensitivity functions, but with a different overall intensity.

#### 4.1.1 Dichromatic system

In large neuronal systems, neurons with similar functional properties can be grouped into homogeneous populations described by their average behavior. Photoreceptors in dichromatic networks, for instance, can be grouped into two main populations characterized by their sensitivity function. Similarly, horizontal cells can be grouped into one or more populations that characterize photoreceptor responses. More specifically, we can think of the network shown in Fig. 1a as a system composed of two populations, *R* and *G*, for red and green photoreceptors, and a single horizontal-cell population, *H*. Chemical synapses and gap junctions between neurons are lumped into effective interactions among populations that are either excitatory or inhibitory, depending on the presynaptic population.

We characterize the average activity of each of these populations by the membrane potential, *h*_*i*_, of the embedded neurons. Currents induced by excitatory and inhibitory synaptic connections are proportional to the number of presynaptic neurons and their corresponding average membrane potential, such that the larger the number of synaptic connections, the stronger the induced internal current. In homogeneous populations, we can consider such internal currents to be a function of the average presynaptic membrane potential, *F*[*h*_*j*_], multiplied by a coupling constant, *w*_*ij*_ [25]. We therefore model the dynamics of these networks using the membrane potentials *h*_*R*_(*λ*), *h*_*G*_(*λ*) and *h*_*H*_(*λ*) of neurons in the corresponding populations, that is,

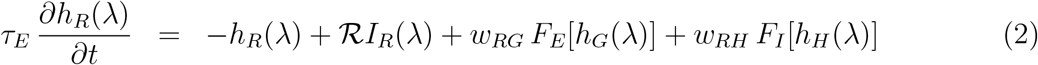

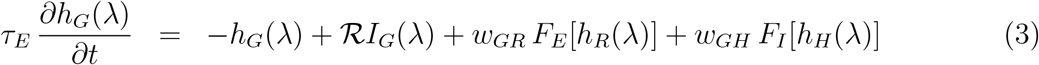

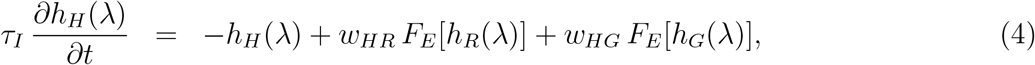

where the last two terms on the right hand sides of Eqs. (2) and (3) correspond to excitatoryand inhibitory connections, respectively, with coupling parameters, *w*_*ij*_, which are positive for excitatory currents and negative otherwise; the first (second) subscript denotes the postsynaptic (presynaptic) population. The functions *F*_*I*_[·] and *F*_*E*_[·], characterize the population response to either inhibitory or excitatory currents, respectively. The second term, *I*_*i*_, corresponds to the independent photoreceptor response described by Eq. (1), and ℛ is the membrane resistance, which we set to unity[25]. Since horizontal cells do not receive direct external stimulation, the only source terms in Eq. (4) correspond to the currents due to photoreceptor populations. The explicit dependence on the wavelength in Eqs. (2)-(4) characterizes chromatically diverse external stimuli; for simplicity, we omit this dependence in all subsequent equations.

The membrane time constants on the left hand side of Eqs. (2)-(4), *τ*_*E*_ for excitatory neurons and *τ*_*I*_ for inhibitory neurons, introduce two time scales that are related to neuron responses and the latency of the feedback mechanism. Some experimental works, reviewed in ref. [12], show the existence of two fast feedback mechanisms from HCs; ephaptic and proton-mediated feedback, highlighting the suitability of HC for tasks involving fast adjustment of cone responses. We take it into consideration, assuming that the time membrane constant of inhibitory neurons is much faster than the time membrane constant of excitatory neurons, *τ*_*I*_ ≪ *τ*_*E*_, which allows us to simplify Eqs. (2)-(4) as,

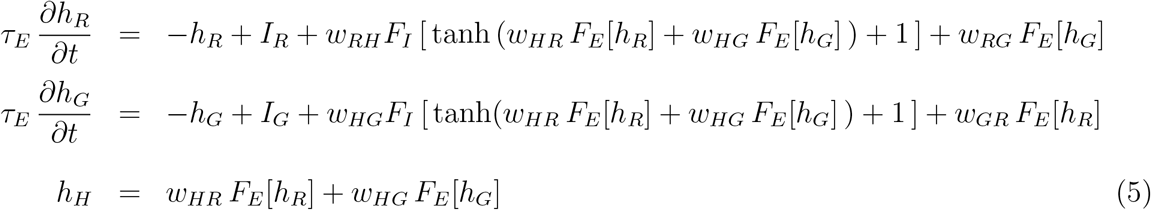

where we have set *F*_*I*_[*h*] = tanh *h*_*h*_ + 1. This response function is always positive and saturates with strong stimulation, resembling typical cone activity. (see Supplementary Material for further details on the response functions).

As previously mentioned, the experimental observations of zebrafish outer retinal layers lead us to ask whether networks of Type I provide an advantage in terms of chromatic information codification and transmission. As noted above, inter-cone gap-junctions have been observed in some vertebrates; the question is whether they are involved in chromatic discrimination. The set of equations (5) provides a simplified framework to study the time evolution of population membrane potentials in dichromatic networks as a function of the synaptic strengths, allowing us to compare Type I networks, with only inhibitory feedback, and Type II networks, with strong inter-cone connections. Comparing the dynamics of the two types of network should provide insights into codification performance.

A Type I network is characterized by weak inter-cone connections, that is, *w*_*RG*_ = *w*_*GR*_ ≈ 0 in Eq. (5). This regime allows an analytic solution of the stationary state as a function of the four remaining coupling parameters, and all the chromatic stimuli illustrated in Fig. 1c. Assuming a saturating excitatory response function, *F*_*E*_[*h*] = tanh(*h*) + 1, We find that for any set of parameters, there is a unique stable fixed point in the two dimensional domain {*h*_*r*_ × *h*_*g*_, ∈ ℝ^2^}. Figure 2a shows a typical phase portrait.

**Figure 2:**
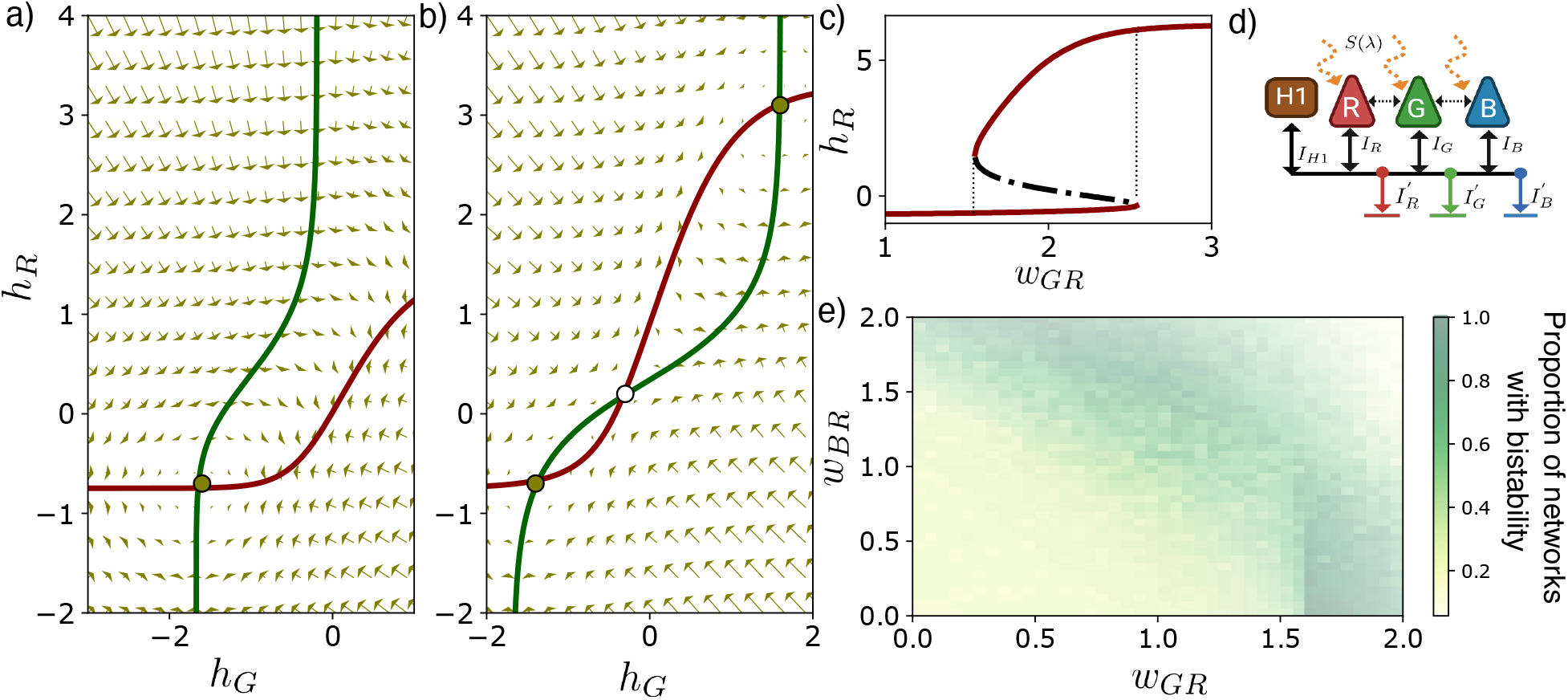
(a, b) Phase portraits of Eq. 5 for a dichromatic network with red and green photoreceptors, and with the parameters *w*_*HR*_ = 1.5, *w*_*RH*_ = −1.7, *w*_*HG*_ = 0.9, *w*_*GH*_ = −1.1, *S*(*λ*) = N (*λ* = 380, 1), and a) *w*_*GR*_ = *w*_*RG*_ = 0 corresponding to Type I, and b) *w*_*GR*_ = *w*_*RG*_ = 1.8, corresponding to a Type II network. c) Bifurcation diagram of *h*_*R*_ for the inter-cone coupling parameters *w*_*GR*_ = *w*_*RG*_ = [1, 3.5]. d) Trichromatic sketch of a fully connected network with an external stimulus, *S*(*λ*). R: red cones, G: green cones, B:blue cones, H: horizontal cells. Solid black arrows represent inhibitory synaptic conections on cones from horizontal cells. Dashed black arrows represent excitatory synaptic connections between cones. e) Proportion of networks exhibiting multistability. Color intensity represents the normalized sum over the discrete interval *w*_*RG*_ = [0.1, …, 2.0], for a fixed combination of the excitatory parameters, *w*_*GR*_ and *w*_*BR*_. All points correspond to the same combination of inhibitory parameters used before.

The existence of a single fixed point implies that the stationary response to a stimulus *S*(*λ*) is unique and independent of the initial state. Our results show that regardless of variations in the coupling strengths among neuronal populations, networks with only inhibitory feedback exhibit a unique response to a given chromatic stimulus, hence reliable codification of chromatic stimuli. In the outermost retinal layers, this is desirable to avoid ambiguity in chromatic codification and transmission.

We now ask whether such behavior persists in a Type II network. We calculate the fixed points of Eqs. (5) by determining the intersections of the nullclines for all coupling parameters in the discrete space *w*_*ij*_ ∈ [0.1, 0.2, …, 5.0) for excitatory and *w*_*ij*_ ∈ (−5.0, …, −0.1, 0) for inhibitory parameters. As shown in Figures 2b-c, we find that, in contrast to Type I networks, Type II networks with strong inter-cone couplings (compared with HC-couplings), can exhibit three fixed points, two of them stable and one saddle node. Under this scenario, a photoreceptor may exhibit multiple responses to the same stimulus, which, as previously mentioned, could lead to misleading or ambiguous chromatic signals. Networks with other photoreceptor combinations, such as red-blue and green-blue, were also studied. As shown in Fig. 3 of the Supplementary Material, such combinations lead to similar phase portraits and dynamical properties, reinforcing our general conclusion.

**Figure 3:**
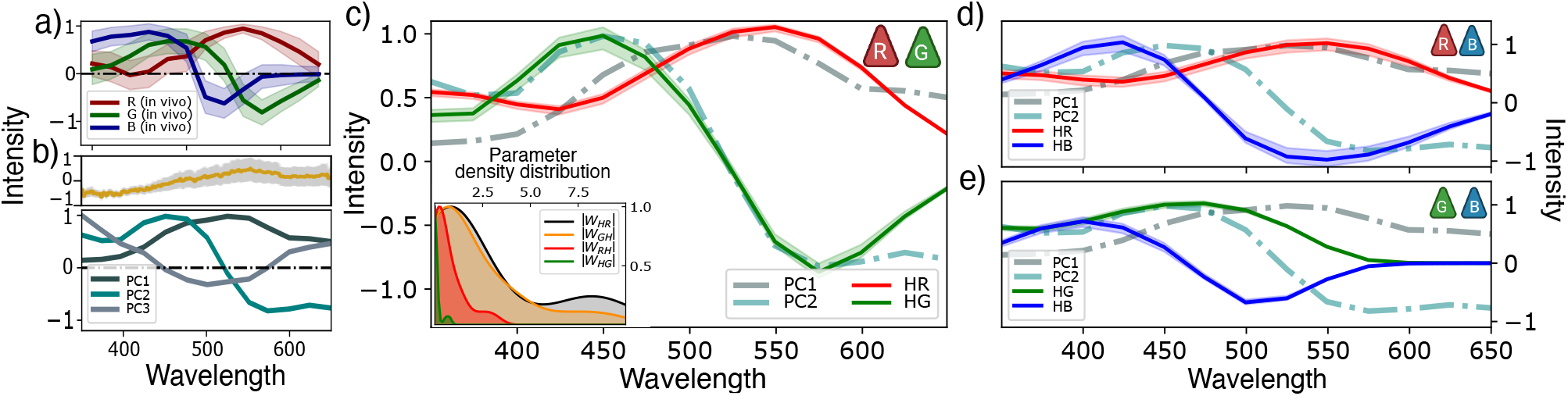
a) In-vivo responses of red, green and blue zebrafish opsins [24]. b) Upper panel: Hyperspectral data from 30 images of aquatic natural images typical of the zebrafish larvae environment [24]. Lower panel: First three principal components obtained from the hyper-spectral data. (c, d, e) Stationary solutions of Eq. (5) for the optimal coupling parameters of a dichromatic network with a) red and green b) red and blue c)green and blue photoreceptors. Inset a) shows the distributions of the optimal parameters (absolute value, |*w*_*ij*_|) over 20 repetitions of the gradient descent algorithm starting from different initial values; inhibitory parameters *w*_*iH*_ are negative by definition; we used a KDE method to infer the curves. Red, green and blue curves correspond to the stationary solutions of the membrane potentials *h*_*r*_, *h*_*g*_ and *h*_*b*_, respectively. Dashed curves correspond to the principal component curves of Fig. 3b.

Thus, we have shown that the responses of a two-photoreceptor system to a given chromatic stimulus are always independent of the initial state of the network if feedback is predominantly inhibitory, via horizontal cells. Conversely, including excitatory inter-cone interactions (e.g., via gap junctions) can lead to responses that depend on the initial state of the network. Experimental observations in zebrafish and drosophila [26] show consistent photoreceptor responses to chromatic stimuli, and the absence of excitatory synapses via gap junctions. The biological reasons why gap junctions among cones are either absent or negligible in these organisms is not fully understood. We find that one explanation for these properties is that they ensure chromatic signal reliability by avoiding multistability and responses dependent on the initial state.

#### 4.1.2 Trichromatic system

The preceding study of networks with two populations of photoreceptors sensitive to different spectral ranges, allows extrapolation of our results to other species with dichromatic retinas. Generalizing these results to species with more complex visual systems, however, is not straightforward. We advance in this direction by investigating trichromatic systems, expanding considerably the diversity of color-vision systems. The trichromatic networks studied here include long- (R), middle- (G) and short-wavelength (B) photoreceptors, and a single horizontal cell population (H) that provides inhibitory feedback to cone populations (see Fig.2d). Assuming that, as in the dichromatic network, *τ*_*I*_ ≪ *τ*_*E*_, the equations of motion are,

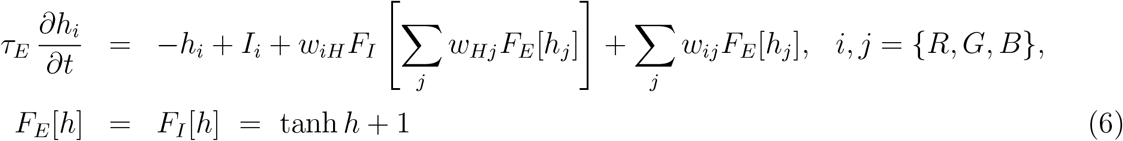

We first investigate whether trichromatic networks with dominant inhibitory feedback (Type I) are also advantageous for chromatic codification, as found in dichromatic networks. Similarly to the dichromatic case, the analytical solution of the system shows that Type I networks exhibit a unique response to the same chromatic stimulus regardless of the coupling parameters strength. To investigate Type-II networks, we characterize the fixed points of Eq. (6) for a discrete set of coupling combinations and for all the chromatic stimuli considered previously. In contrast to dichromatic networks, such fixed points, if any, are embedded in a three-dimensional phase portrait. To locate the fixed points, we minimize a cost function, ℒ, that is zero if and only if 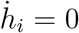 for all three photoreceptor populations in Eq. 6, that is,

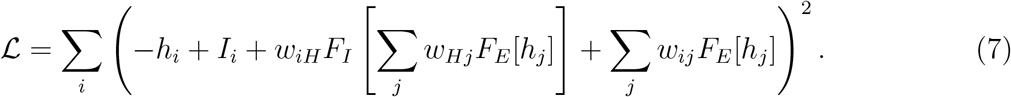

We use a gradient-descent algorithm to find the global minima of ℒ for each parameter combination. It is easy to verify that, depending on the parameter combination, the network can exhibit one, two or three fixed points. Similarly to the dichromatic case, multiple fixed points are common in Type II networks with strong inter-cone connections. To see this in detail, note that Eq. 7 has six excitatory parameters corresponding to the couplings between red, green and blue cone populations. Considering a symmetric interaction between populations, only three free parameters, *w*_*RG*_ = *w*_*GR*_, *w*_*RB*_ = *w*_*BR*_ and *w*_*GB*_ = *w*_*BG*_, remain. For each combination of inhibitory couplings previously studied, we calculate the number of fixed points for all combinations of these three remaining excitatory couplings, finding that the stronger the excitatory parameters, the larger the likelihood of bistability. We summarize this result in the intensity plot of Fig. 2e. For each fixed combination of the parameters *w*_*GB*_ and *w*_*RB*_, we count the number of networks exhibiting multiple stable fixed points when varying the discrete values *w*_*RG*_ = [0.1, 0.2 … 2]. We find that networks with at least one strong excitatory inter-cone coupling tend to exhibit multiple responses to the same chromatic stimulus. This behaviour is similar for all Gaussian stimuli studied.

We conclude that adding a third type of photoreceptor to the outermost retinal networks does not change our general conclusion regarding population interactions. In contrast, these results support the hypothesis that networks with predominantly inhibitory feedback provide an advantage for reliable and unambiguous chromatic codification. As a complementary study, we investigated six other species with different sets of opsins (see Supplementary Fig. 4), finding similar dynamical properties. This suggests that our findings can be generalized to other species.

### 4.2 Optimal chromatic codification

In the previous section, we focused on the synaptic strengths between populations that yield stable responses. In this section, we use both zebrafish in-vivo photoreceptor activities [17] and hyperspectral data from zebrafish environments [8] to investigate the architecture of outer retinal networks from the viewpoint of optimal codification and transmission of chromatic information.

Fig. 3a shows the *in-vivo* spectral responses of zebrafish photoreceptors. Remarkably, interactions with horizontal cells cause green and blue neurons to exhibit opponent responses to chromatic stimuli, which as discussed above, suggests early optimization of chromatic information transmission. Adopting this optimization hypothesis, we expect: (1) Reduced redundancy of network responses in spectral space[11]; (2) Optimal codification of the environmentally available chromatic information. As a proxy for optimal chromatic codification, we use a principal component analysis of the hyperspectral data of aquatic naturalistic images typical of zebrafish environments studied in Ref. [8]. In accord with the analysis of ref. [17], we find that the first three principal components capture more than 97% of the chromatic data variance. Figure 3b shows these three principal components (PC1-PC3), the first without zero-crossings, the second with one zero-crossing, and the third with two. Comparing Figs. 3a and 3b, we observe that the in-vivo red and green cone responses match the first two principal components qualitatively, supporting the hypothesis of optimal codification of chromatic stimuli in zebrafish outermost retinal layers.

We begin our analysis by studying the dichromatic network shown in Fig. 1a and described by Eq. (5). We adjust the coupling parameters, *w*_*ij*_, such that the spectral responses of the photoreceptor populations, *h*_*R*_ and *h*_*G*_, match the first two principal components, which together explain more than 91% of the hyperspectral data variance. (We use the L-BFGS-B minimization algorithm [nnn]to obtain the coupling parameters yielding the best match). Figure 3c shows the solutions of Eq. 5 for a network with only inhibitory feedback (in red and green), with both a no zero-crossing and a single zero-crossing curve, as expected for color-opponent signals[11, 27]. The inset shows the probability densities of the coupling parameter absolute value |*w*_*ij*_| of Eq. 5 over different basins of attraction. Similarly, we adjust the parameters of a network with both inhibitory feedback and excitatory inter-cone connections; in all cases, the optimal excitatory couplings are weak or negligible, leading to results similar to those found for the inhibitory network.

As expected from our previous dynamical analysis, we find a unique stationary fixed point for all optimal networks that exhibit the expected opponent chromatic responses. This means that the membrane-potential dynamics is the same (at least for *τ*_*I*_ ≪ *τ*_*E*_) for any initial condition, allowing reliable information codification of chromatic stimuli. We also studied other opsin combinations, leading to the results shown in figures 3(d, e). We observe that having only blue and red photoreceptors provides a qualitatively good fit to the first principal component, but not to the second. Moreover, dichromatic networks with only blue and green photoreceptors cannot fit either of the first two principal components. Contrasting the results for these two-photoreceptor combinations, we conclude that dichromatic networks with long -and middle-range photoreceptors have the best performance when codifying chromatic information typical of the zebrafish environment.

Next, we analyze trichromatic networks, restricting the parameter space to reproduce the expected opponent responses, as we have done for dichromatic networks. Following the same procedure as before, we fit the membrane potential response of each photoreceptor population, *h*_*R*_, *h*_*G*_ and *h*_*B*_, to the first three principal components of the naturalistic images. As shown in Fig. 4a, the optimized network yields a poor fit to the third principal component, and the network responses do not match qualitatively the expected optimal responses. Instead, we find a single zero-crossing in the blue photoreceptor response curve. We note, though, that in-vivo recordings of photoreceptor responses do not match this third principal component either, as shown in Fig.3a.

**Figure 4:**
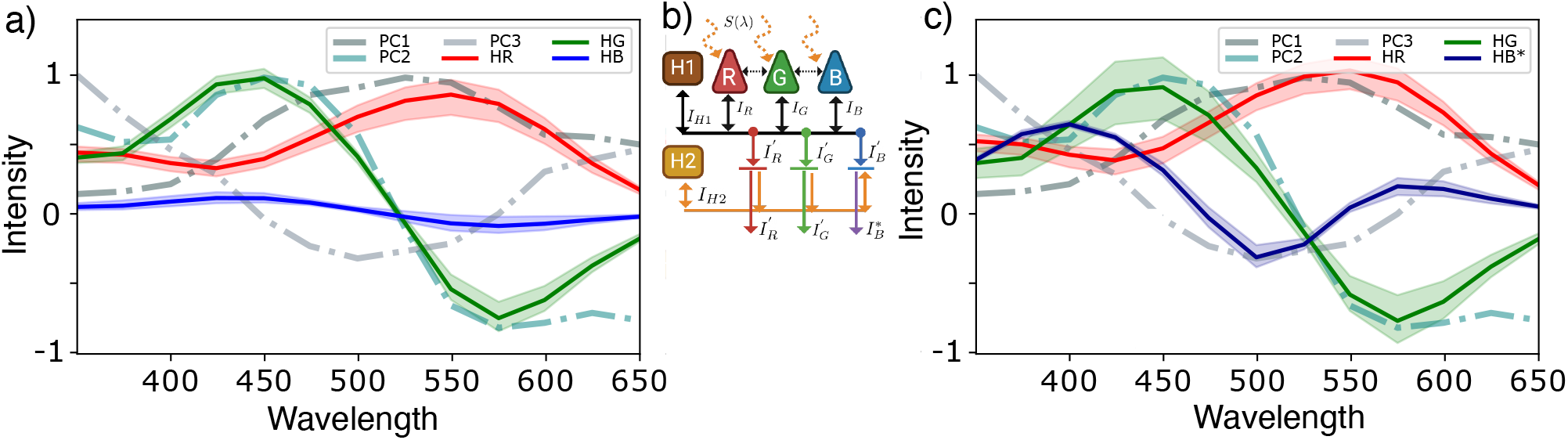
a) Stationary solutions of Eq. (6) for the optimal coupling parameters of the trichromatic network sketched in the inset. b) Sketch of a fully connected trichromatic network with two types of horizontal cells providing two successive inhibitory feedback mechanisms. c) Stationary solutions of Eq. (6, 8) for the optimal coupling parameters of the trichromatic network sketched in the inset. Red, green and blue curves correspond to the stationary solutions of the membrane potentials *h*_*r*_, *h*_*g*_ and *h*_*b*_, respectively. Dashed curves correspond to the principal component curves of Fig. 3b.

For trichromatic systems, type I and type II networks are unable to reproduce all three principal components, leading as to ask whether us to ask whether an expanded network, e.g., with a second parallel inhibitory feedback or with a fourth type of photoreceptor, is capable of realizing this task. We begin by including a second type of horizontal cell, H2, providing feedback to only two of the three cone populations. Repeating the previous analyses, we find that networks with two parallel inhibitory feedback mechanisms do not eliminate the errors in fitting of the third principal component. Similarly, we investigated whether a fourth type of photoreceptor, with a maximum response in the UV range, might allow the other three photoreceptor responses to match all three principal components simultaneously. We found that adding such a UV cone population has little effect on the functional responses of the other cones, in agreement with the experimental analysis of zebrafish by Yoshimatsu et al.[17]. Details of these analyses are provided in the Supplementary Material.

From these results we conclude that in zebrafish retinas, trichromatic networks are incompatible with obtaining the first three principal components of environmental chromatic data at the first synaptic connection. This motivates us to add a second synaptic connection, or network layer, to verify whether such a network can be optimized to reproduce the expected results. Figure 4b shows a sketch of such a network, with a second type of horizontal cell, H2*, integrating the first layer responses, and providing feedback only to population B, which is the one unable to fit the third principal component. The blue cone population response, 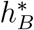, at the second layer is described by equation,

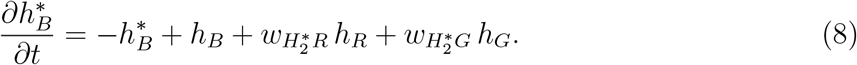

with 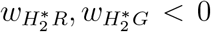. Figure 4c shows the responses of the optimal network after learning the parameters of equations 6 and 8. The improved agreement between response curves and PCs suggests that obtaining the three optimal chromatic channels requires a second synaptic contact (or network layer), including a second type of inhibitory neurons. Although we modeled this population as a second type of horizontal neuron, other neuronal populations in inner retinal layers, such as bipolar cells, might also realize this task. We also stress that the first three principal components explain more than 97% of the hyperspectral data variance, but that the third PC alone explains less than 6%. Consequently, despite the fact trichromatic networks with a unique inhibitory feedback mechanism do not reproduce the expected third principal component, the other two channels capture more than 91.3% of the hyperspectral data variance, which is still a good performance.

In summary, adopting ideas from information theory and chromatic opponency, we have shown that after the first synaptic connection, dichromatic networks can yield optimal responses when red cones are available. Otherwise, responses of photoreceptor populations are suboptimal. Generalizing to trichromatic networks, we find that, after the first synaptic connection, photoreceptors are unable to exhibit all three opponent responses. Only by including a second synaptic connection, or network layer, with a fourth neuronal population, can photoreceptor responses match the PCs.

#### 4.2.1 Optimal sensitivity curves

Our study identified networks that optimize the codification of chromatic information available in the zebrafish environment, using as a template of photoreceptor responses the set of zebrafish sensitivity curves. Now we ask whether the zebrafish opsins are optimal at fitting the given chromatic information, or if instead, there are other opsin combinations leading to more precise chromatic codification. Altough optimization is not the only feature that determines visual system properties, it has been shown that some species adapt aspects, such as the sensitivity functions, to fit the environmental conditions[28, 29].

To answer this question, we contrast the performance of networks composed of different opsin combinations obtained by varying the wavelength of maximum sensitivity of zebrafish cones, while maintaining the shape of the distribution fixed (we use a discrete step interval of Δ_*λ*_ ≈ 12nm). For all opsin combinations, we calculate the optimal two-layer network, described by Eqs. (6) and (8), fitting the first three principal components of the zebrafish environment, as in the previous sections. To quantify the network performance, we define the cost function,

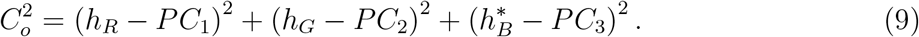

Using the same invertal Δ_*λ*_ for blue and green opsins, we calculate the cost function forall possible combinations within the interval (350, 650)nm, and search for an optimal set of opsins. Figure 5a shows the intensity plot of the cost function for all green and blue opsin combinations given a fixed red sensitivity curve. We find the global minimum is reached at 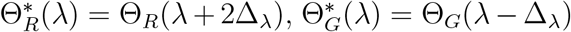 and 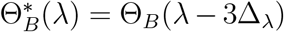, as illustrated in Fig. 5b,c.

**Figure 5:**
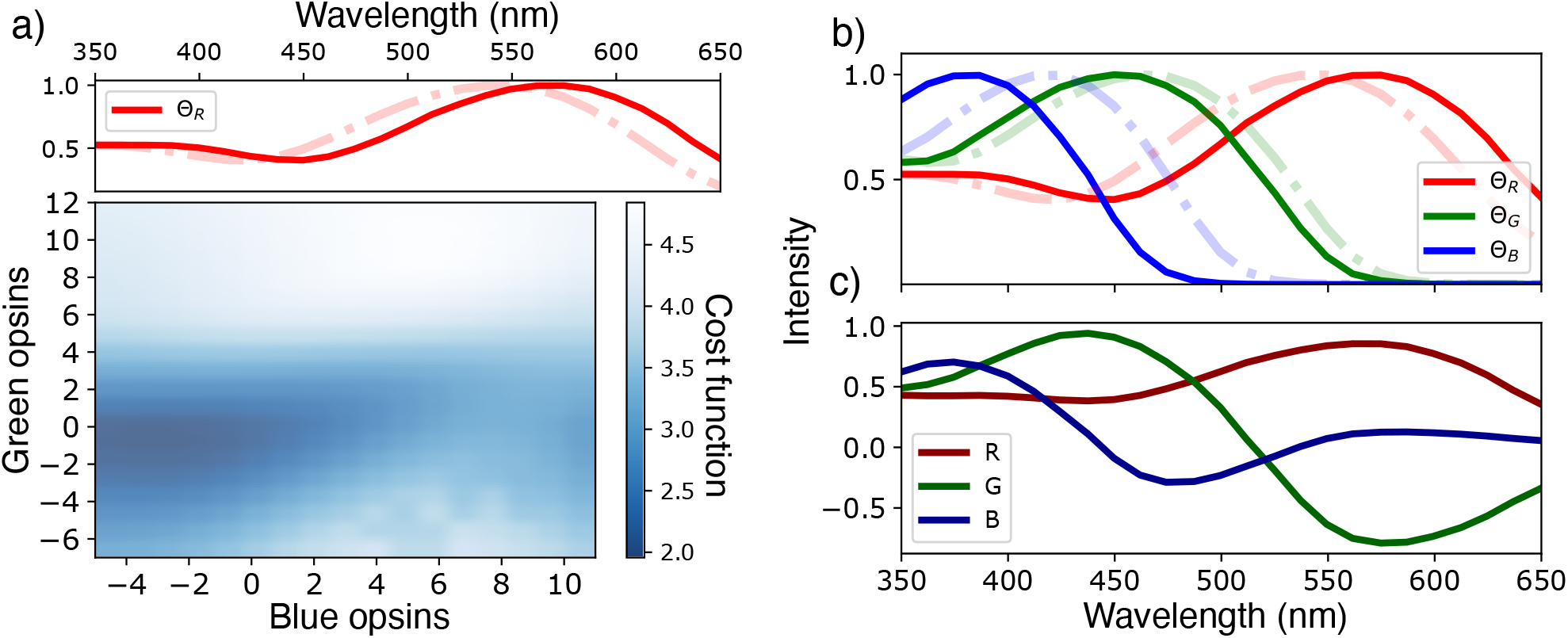
a) Intensity plot of the cost function (Eq. 9) for different opsin curves combinations, darker colors represent smaller values. Red opsin was first fixed to optimize all possible combinations; the optimal curve is shown in the upper plot. X and y labels in the intensity plot represent the number of shifts (in steps of 5nm) of blue and green curves respectively, with the sign indicating the shift direction. b) Comparison between experimentally observed sensitivity functions (dashed lines) and the optimal fitting curves (solid lines). Arrows indicate the optimal shift direction. c) Optimal network responses for this optimal set of opins.

We conclude that compared with zebrafish, the optimal set of cones improves performance by approximately 13%, suggesting that retinal networks with zebrafish cones are near to optimal codification. Extrapolating these results to other species with similar environmental chromatic conditions requires knowledge of the independent cone responses, or sensitivity curves to properly compare the networks. Studies involving species with different environmental conditions would naturally require analysis of the corresponding environmental hyperspectral data.

## 5 Discussion

Identifying the retinal circuits specialized in chromatic discrimination leads to an understanding of how organisms extract spectral information from their environment[2, 30]. In-vivo experiments on outer retinal circuits, though, are only feasible in a restricted number of species, limiting a broader study. In this situation, theoretical models are useful to predict general features that can be tested in such model organisms and extrapolated to others beyond experimental access. Our study attempts to identify features of zebrafish-inspired outer retinal networks to understand more broadly the biological fundamentals of chromatic codification and transmission in visual systems. As mentioned earlier, we focused our work on networks that mediate chromatic information via HCs. Other species, such as butterflies [23], with different network architectures will be studied in future works.

In the first part of our study, we found that in outer retinal networks with fast inhibitory feedback, inter-cone excitatory connections can lead to bistability, so that network responses to a given chromatic stimulus could be ambiguous. Ideally, one expects photoreceptors to codify as much relevant visual information as possible from the environment and transmit it reliably to downstream circuits. More complex tasks, such as perception or recognition, for which such bistability might be desirable, are supposed to take place later in the visual system. Our results suggest that, in retinas, inter-cone gap junctions, if present, are likely not involved in chromatic codification. Some experimental studies on macaques and other vertebrate species [20, 31, 32] have shown evidence of both cone-cone and cone-rod gap junctions in the foveal region. The role of such connections, however is non fully understood. In ground squirrel retinas, for instance, such connections seem to play an important role in increasing the signal-to-noise ratio [31]; similar behaviour has been found in macaque retinas [21]. Noise reduction is highly relevant in low-luminance environments. At high luminance, where color discrimination is possible, the effects of gap junctions seem to be negligible, supporting the hypothesis that gap junctions are involved in achromatic tasks. Some other works [33] have shown evidence of excitatory feedback in individual synapses between HCs and cones, which might lead to other effective connections, not considered in our model.

In addition to studying different types of feedback, we investigated network architectures leading to optimal codification of the available chromatic information. Determining whether outer retinal circuits are optimized to codify available chromatic information allows one to gauge the relevance of color discrimination in a given species. Such findings might afford insights into other areas, such as ecology and animal behavior. As suggested by Yosimatsu et al. [17], zebrafish seem to codify of the chromatic information efficiently, qualitatively matching the first two PCs of the hyperspectral data. With our model, we find similar results. Specifically, we found that zebrafish-inspired outer retinal networks with a single HC layer can reproduce only the first two PCs of the zebrafish hypterspectral data, explaining 91% of the variance. Capturing more than 97% of the variance requires fitting all three PCs. We find that a more complex network, with two interneuron layers is necessary. We would expect retinas to optimize the trade-off between information transmission and metabolic cost, such that information transmission is guaranteed at the lowest energy expenditure. Nevertheless, even with an optimal architecture, more complex networks are necessary to transmit specific information, as suggested by our results. Whether it is advantageous for an organism to invest energy increasing an additional retinal layer to improve chromatic discrimination depends on species-specific interactions with its habitat. These results are general for networks with inhibitory interneuronal layers and similar environmental conditions.

Our study of network architecture was performed using zebrafish opsin curves as a reference. Subsequently, we investigated whether other opsin combinations might lead to further optimization of the network, that is, whether it would be possible to improve zebrafish performance at codifying environmental chromatic information. We find that optimizing opsin combinations leads to an improvement of approximately 13% in zebrafish performance. As mentioned earlier, in some species, such as bees, photoreceptor sensitivity functions are highly tuned to the environmental chromatic spectrum, leading to an optimal codification of color information[28, 29]. Our results on zebrafish show that they have close-to-optimal chromatic codification, but that there are other photoreceptor combinations yielding modest improvements. Other relevant features, such as movement or edge detection, might favor retinal circuits that optimize other general aspects of visual stimuli, by penalizing slightly chromatic information. As before, these results hold for zebrafish-inspired retinal circuits under similar environmental conditions.

## Acknowledgments

We thank Tom Baden and his group for helpful discussions and for sharing the experimental data. We acknowledge helpful discussions with L. Abbot, R Martinez-Garcia, X Chen and M Bauer. Work supported by the Consejo Nacional de Desarrollo Cientifico y Tecnologico, CNPq.

## Notes

### Competing Interest Statement

The authors have declared no competing interest.

