## Supplementary Material for "Data-driven models of optimal chromatic coding in the outer retina"

#### 1 Fixed points in dichromatic networks

In the stationary state, Eq. (5) (main text) for Type I networks leads to the nullcline equation,

$$h_r^* = \frac{w_{RH}\, h_G^* + w_{GH}\, I_R - w_{RH}\, I_G}{w_{GH}}, \tag{1}$$

showing that for a network with a given set of coupling parameters $w_{ij}$, there is a unique fixed point, $\mathbf{h}^* = \{h_R^*, h_G^*\}$.

For type II networks, we get the nullcline equations,

$$\begin{aligned} w_{GH}\, h_R + w_{RH}\, w_{GR}\, \tanh h_R &= w_{RH} h_G + w_{GH} w_{RG} \tanh h_G + w_{GH}\, I_R \\ &\quad -w_{RH}\, I_G + w_{RG}(w_{GH} - w_{RH}). \end{aligned} \tag{2}$$

We calculate the fixed points in the space of the coupling parameters, as shown in Figs. 2(a, b) (main text). We determine the stability of the fixed points via the Jacobian matrix, that is,

$$
\begin{aligned}
J &= \begin{pmatrix} -1 + w_{RH}\, w_{HR}\, \mathrm{sech}^2\, h_R\, \mathrm{sech}^2\, D & \mathrm{sech}^2\, h_G (w_{RH}\, w_{HG}\, \mathrm{sech}^2\, D + w_{RG}) \\ \mathrm{sech}^2\, h_R (w_{GH}\, w_{HR}\, \mathrm{sech}^2\, D + w_{GR}) & -1 + w_{GH}\, w_{HG}\, \mathrm{sech}^2\, h_G\, \mathrm{sech}^2\, D. \end{pmatrix} \\
&= \begin{pmatrix} J_{11}^{(1)} + J_{11}^{(2)} & J_{12}^{(1)} + J_{12}^{(2)} \\ J_{21}^{(1)} + J_{21}^{(2)} & J_{22}^{(1)} + J_{22}^{(2)} \end{pmatrix} \qquad (3) \\
D &= \tanh(w_{HR}\, F_E[h_R] + w_{HG}\, F_E[h_G]) + 1
\end{aligned}
$$

The trace of this matrix is negative for all coupling parameters. On the other hand, the determinant sign depends on the coupling parameter strengths. If Det $< 0$ the fixed point is classified as a saddle point. Otherwise, it is a stable sink.

For trichromatic networks, we use a gradient-descent algorithm to find the nullcline intersection points using Eq. (7) (main text), and similarly, we determine the stability of the fixed points using the corresponding Jacobian matrix.

### 2 Response functions

Photoreceptor responses to external stimuli increase up to a certain threshold, at which they saturate. We model this behavior using the functions $F_E[h] = F_I[h] = \tanh h + 1$, as shown in Fig. 1 (solid line). Strong hyperpolarized (depolarized) regimes of the pre-synaptic neuronal population, induce negligible (significant) post-synaptic responses.

Depending on the specific neuron current integration, it is possible to further specify the response using the more general function $F[h] = \tanh(\alpha h + \beta) + 1$, whose parameters might vary for both excitatory and inhibitory pre-synaptic populations (see Fig. 1). We investigated the stability of this new system by comparing the Jacobian with Eq. of the supplementary material, that is,

$$J = \begin{pmatrix} J_{11}^{(1)} + \alpha_I\,\alpha_E\,J_{11}^{(2)} & \alpha_I\,\alpha_E\,J_{12}^{(1)} + \alpha_E J_{12}^{(2)} \\ \alpha_I\,\alpha_E\,J_{21}^{(1)} + \alpha_E J_{21}^{(2)} & J_{22}^{(1)} + \alpha_I\,\alpha_E\,J_{22}^{(2)}. \end{pmatrix} \tag{4}$$

This means that because $\alpha_E = \alpha_I > 0$, the stability portraits discussed in the main text remain equal, keeping our general results unchanged.

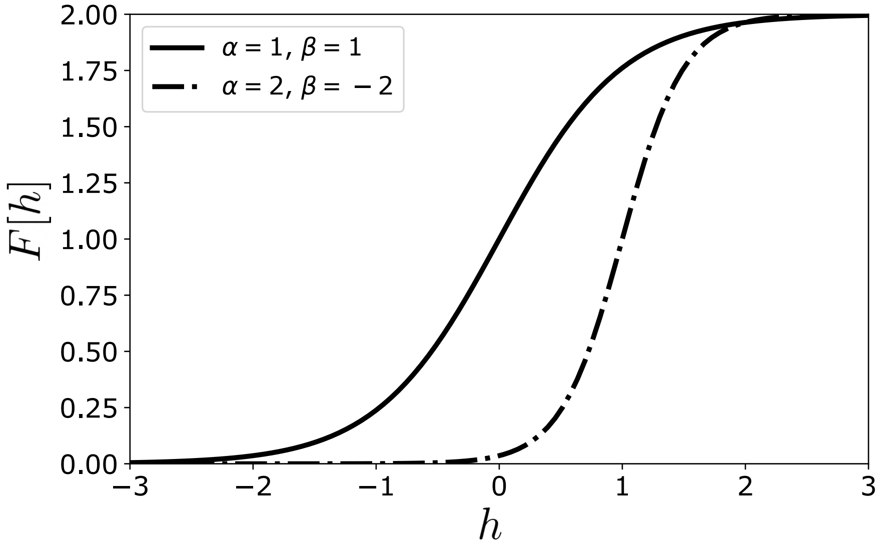


Figure 1: Response functions for the parameters $\alpha = \beta = 1$ (solid line) and $\alpha = 2, \beta = -2$ (dashed line).

### 3 Generalized network: UV photoreceptors and multiple horizontal cells

To study whether a fourth photoreceptor type would improve the network response, we include an ultraviolet photoreceptor in our formalism (see Fig. 2a). As shown in Fig. 2b, the responses of both the functional and isolated UV cones are similar, suggesting the absence of significant feedback from other populations to UV cones. We include a new population (U) that feeds the HC population (H), but does not receive excitatory/inhibitory feedback. The equation of motion remains the same as Eq. (6), but now with $i, j \in \{R, G, B, U\}$, and $W_{Ui} = W_{UH} = 0$. As shown in Fig. 2c, the gradient-descent algorithm to fit the coupling parameters yields a poor fit to the third principal component, as in the trichromatic network studied in the main text. We also included unilateral excitatory connections $W_{Ui}$, finding a maximum improvement of $< 2\%$ in the blue photoreceptor response. This suggests that ultraviolet cones do not play a key role in the chromatic opponent process of red, blue and green zebrafish photoreceptors.

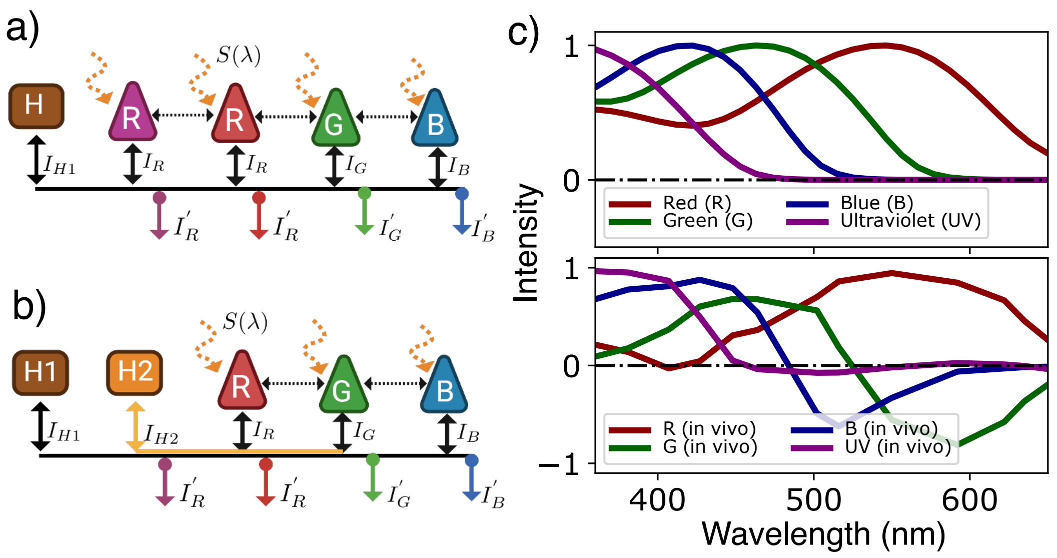


Figure 2: Tetrachromatic network: a) sketch of a network with an external stimulus, $S(\lambda)$. R: red cones, G: green cones, B:blue cones, U: uv cones, H: horizontal cells. Solid black arrows represent inhibitory synaptic conections between cones and horizontal cells. Dashed black arrows represent excitatory synaptic connections between cones. b) Sensitivity functions of independent red, green, blue and uv zebrafish opsins. d) In-vivo responses of red, green, blue and uv zebrafish opsins. c) Stationary solutions for the optimal coupling parameters of the tetrachromatic network sketched in a). Red, green and blue curves correspond to the stationary solutions of the membrane potentials $h_r$, $h_g$ and $h_b$, respectively. Dashed curves correspond to the principal component curves of Fig. 3b. d) Trichromatic network sketch of a fully connected network with an external stimulus, $S(\lambda)$, and two inhibitory feedback mechanisms. R: red cones, G: green cones, B:blue cones, H1: first population of horizontal cells, H2: second population of horizontal cells. Solid black arrows represent inhibitory synaptic conections between cones and horizontal cells. Dashed black arrows represent excitatory synaptic connections between cones.

In addition, we studied a trichromatic network with two fast-parallel inhibitory mechanisms: the first integrating responses from all three cone populations and the second integrating responses from only two cone populations (see Fig. 2d). The dynamical equations are,

$$\tau_E \frac{\partial h_i}{\partial t} = -h_i + I_i + \sum_{j \neq i} (w_{ij} F_E[h_j]) + w_{iH_1} F_I \left[ \sum_j w_{H_1 j} F_E[h_j] \right] + w_{iH_2} F_I \left[ \sum_{j'} w_{H_2 j'} F_E[h_{j'}] \right]$$

$$h_{H_1} = w_{H_1 R} F_E[h_R] + w_{H_1 G} F_E[h_G] + w_{H_1 B} F_E[h_B]$$

$$h_{H_2} = w_{H_2 R} F_E[h_R] + w_{H_2 G} F_E[h_G].$$

The terms including the second inhibitory feedback mechanism did not change significantly the performance of the network to fit the principal component curves.

### 4 Supplementary Figures

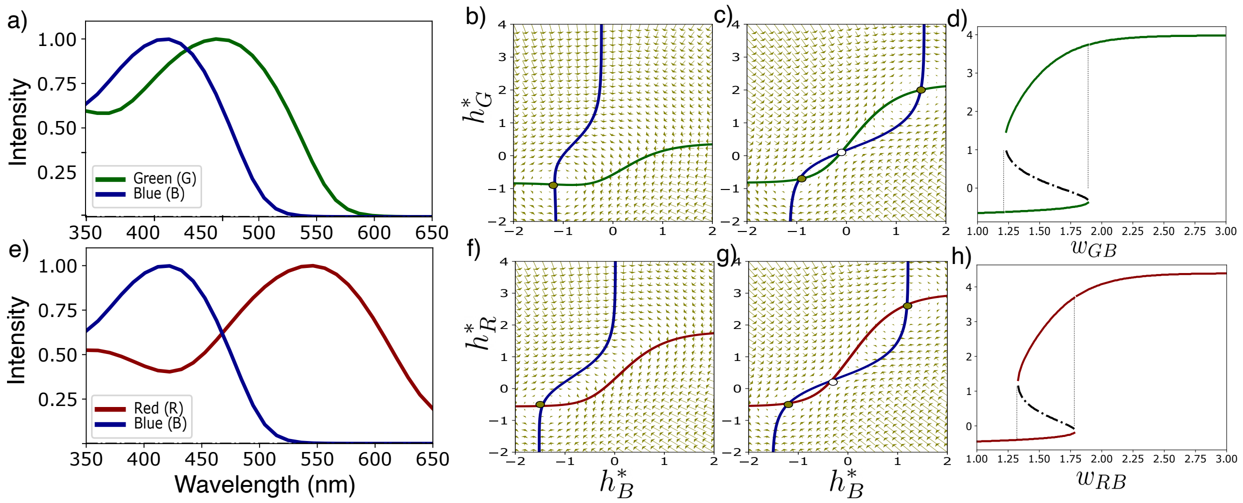


Figure 3: a) Green and blue zebrafish photoreceptor sensitivity curves. (b, c) Phase portraits of Eq. (5) for a dichromatic network with green and blue photoreceptors, and with the parameters a) $w_{HG} = 0.9, w_{GH} = -1.1$, $w_{HB} = 1.5$, $w_{BH} = -1.5, w_{BG} = 0$ and $S(\lambda) = \mathcal{N}(\lambda = 412, 1)$, corresponding to a Type I network, and b) $w_{HG} = 0.9, w_{GH} = -1.1$, $w_{HB} = 1.5$, $w_{BH} = -1.5, w_{BG} = 1.6$ and $S(\lambda) = \mathcal{N}(\lambda = 412, 1)$, corresponding to a Type II network. d) Bifurcation diagram as a function of the coupling parameter $w_{GB} = w_{BG}$. e) Red and blue zebrafish sensitivity curves. (f, g) Phase portraits of Eq. (5) for a dichromatic network with red and blue photoreceptors, with parameters a) $w_{HR} = 0.9$, $w_{RH} = -1.1$, $w_{HB} = 1.5$, $w_{BH} = -1.7$, $w_{RB} = 0$ and $S(\lambda) = \mathcal{N}(\lambda = 390, 1)$, corresponding to a Type I network, and b) $w_{HR} = 0.9$, $w_{RH} = -1.1$, $w_{HB} = 1.5$, $w_{BH} = -1.7$, $w_{RB} = 1.7$ and $S(\lambda) = \mathcal{N}(\lambda = 390, 1)$, corresponding to a Type II network. h) Bifurcation diagram as a function of the coupling parameter $w_{RB} = w_{BR}$.

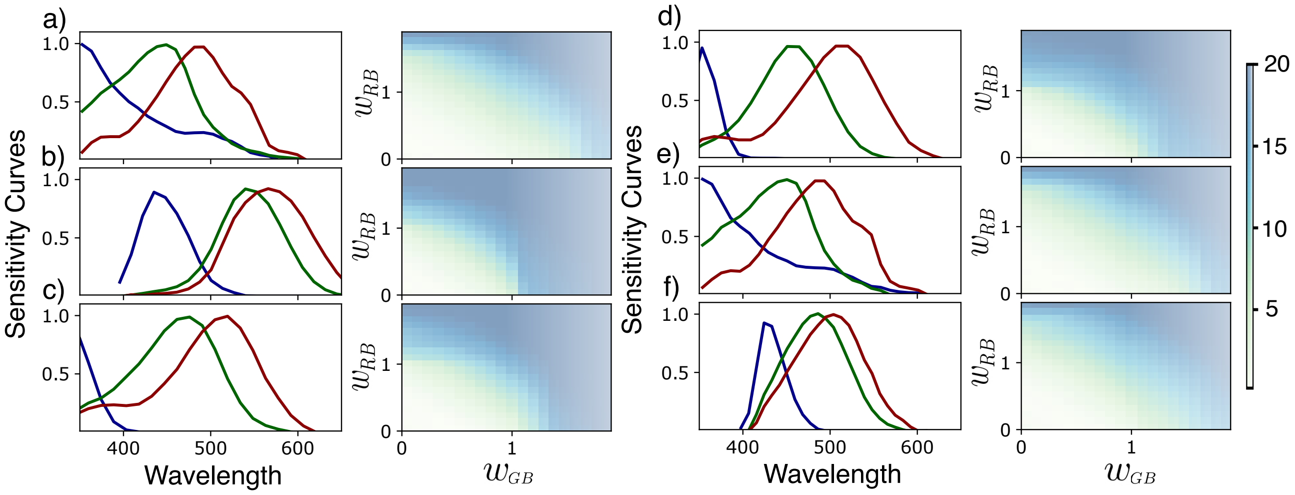


Figure 4: Sensitivity curves (left) and number of networks with multiple fixed points when varying $w_{RG} = [0, 0.1, \ldots, 2.0)$, for the fixed parameters $W_{rh} = W_{hg} = 0.1$, $W_{hr} = W_{bh} = 0.2$ and $W_{gh} = W_{hb} = 0.3$, for: (a) Honeybee. (b) Human. (c) Spider. (d) Damselfish. (e) Giant clam. (f) Triggerfish. Data on sensitivity curves were extracted from Fig. 7.2 in reference [3].
